# Unfolding of the disease-linked bank vole prion protein E200K in the context of PrP^Sc^ propagation

**DOI:** 10.64898/2026.10.13.759008

**Authors:** Sanaz Sabzehei, Hasier Eraña, Iria Díaz-Arias, Raúl Cacheiro, Arcadio Guerra, Sara María Álvarez-Cardama, Yaiza de León Vega, Human Rezaei, Joaquín Castilla, Víctor M. Sánchez-Pedregal, Jesús R. Requena, Manuel Martín-Pastor

## Abstract

The elucidation of the atomistic structure of PrP^Sc^ has, disappointingly, not revealed the mechanism underlying prion propagation. This is because PrP^Sc^ has been shown to be a standard amyloid, and as such, it can only template a polypeptide chain that is unfolded. Therefore, while PrP^Sc^ can easily template the disordered ∼90-120 domain of an incoming PrP^C^ molecule, it does not have any obvious capacity to template its ∼121-231 folded domain (FD). The FD needs to accommodate into the ∼121-230 PrP^Sc^ surface, an inert “Procrustean bed”. We have recently proposed that unfolding of the FD is a prerequisite for its conversion to the PrP^Sc^ conformation and using experimental and computational methods identified the likely early events of such process using bank vole (BV)PrP(90-231) as our working model. We concluded that a key early event is the destabilization of the short β1-β2 assembly, and that the segment contiguous to the disordered tail, ∼121-140, encompassing β1 and its adjacent coils, is the most likely region to unfold first. In the present study we applied the same methodology to the disease-related PrP mutant E200K. E200K is the most prevalent mutation leading to familial prion disease in humans, yet the stability of its FD is only marginally lower compared to that of the protein with the wild type (wt) sequence. We, therefore, aimed at assessing whether the early unfolding events of BVPrP^C^(E200K) are similar or different to those of wtBVPrP^C^ and whether such early unfolding route might provide insight into the extreme capacity of PrP^C^(E200K) to spontaneously generate PrP^Sc^. It should be noted that generation and propagation of PrP^Sc^ are distinct phenomena, even though both are mechanistically related and dependent on conformational changes in PrP^C^. Our data show that early unfolding events of BVPrP^C^(E200K) are essentially similar to those of wt BVPrP^C^. However, interesting nuances exist.

## Introduction

It might have been believed that elucidation of the atomistic structure of PrP^Sc^ would lead to an immediate understanding of the mechanism of prion propagation (for reviews on these unique protein-only pathogens, see (1-3). However, PrP^Sc^ is just a parallel-in-register beta stack (PIRIBS)-based amyloid (4-8), and as such it can only template a previously unfolded polypeptide chain, a capacity of all amyloids (9-11). Therefore, while conceivably PrP^Sc^ can easily template the disordered ∼90-120 domain of an incoming PrP^C^ molecule, it cannot directly template its ∼121-231 FD (bank vole numbering). We have recently proposed that the FD needs to unfold first to accommodate into the ∼121-230 PrP^Sc^ surface, an inert “Procrustean bed” (12-13).

We proposed that once the disordered ∼90-120 segment of PrP^C^ has adopted the PrP^Sc^ conformation, the remaining globular FD (residues ∼121-230), that comprises three α-helices, two short antiparallel β-strands, and several connecting loops (14-15) (Supplementary Fig. S1), remains tethered in front of the PrP^Sc^ templating surface. Over time, breathing movements within the FD should lead to temporary unfolding of some short segments, which could be trapped by their isosequential tracts on PrP^Sc^, acting as a ratchet mechanism. Using thermal unfolding and NMR spectroscopy, we identified the ∼121-140 segment as the most likely region to unfold first, with β1 as a very relevant hotspot, based on high temperature coefficient (Tc) values, accumulation of non-linear CCSD values, other NMR-derived parameters and molecular dynamics (12). Physical proximity of this region to its docking area on the PrP^Sc^ surface is also relevant.

A corollary of our proposal is that PrP^C^ molecules with a less stable FD should unfold more easily and therefore, in general, convert to the PrP^Sc^ conformation (in the presence of templating PrP^Sc^) more readily. In support of this, Kong *et al*. generated a transgenic mouse line expressing a human PrP with the V209M mutation [HuPrP(V209M)], which exhibits increased thermodynamic stability compared to wtHuPrP. When these mice were challenged with (wt) sCJDMM1 prions, they showed incomplete attack rate with very long incubation times. This could be explained as a consequence of the transmission barrier created by the mismatch between the wt template (wtPrP^Sc^ inoculum) and the mutation in the PrP^C^ substrate. However, upon abrogation of this effect through second and third passages, incubation times were still longer in the mutant mice expressing a less unfoldable PrP^C^ with respect to mice expressing the same level of wtHuPrP (16).

On the other hand, it is noteworthy that while a few disease-related mutations result in PrP^C^ species with decreased stability, like T183A (17), many others do not confer decreased stability (18). Furthermore, these mutant PrP^C^ sequences must have an increased tendency to unfold in a way that results in generation *de novo* of a first pool of mtPrP^Sc^. This propensity might in many cases be related to subtle structural characteristics of the mutant PrP^C^ molecule leading to specific unfolding pathways that are relevant to association with other unfolding PrP^C^ units In a way that yields an initial PrP^Sc^ architecture, but not necessarily to an increased tendency to unfold in the context of PrP^Sc^-templated propagation.

An interesting case is the E200K mutation. E200K is the most prevalent disease-related mutation in humans (19, 20). The solution structure of this mutant HuPrP^C^ FD is essentially undistinguishable from that of wtHuPrP^C^ (15). Its stability is only slightly lower than that of the wt protein, with a Tm of 67.0 ± 0.9 °C for HuPrP^C^E200K(90-231) *vs*. 70.2 ± 0.1 °C of the wt (21). On the other hand, the mutation results in a notable perturbation of the electrostatic potential (15) and a loosening of the structure of helix α3 as a consequence of the disappearance of the salt bridge between residues E200 and K204 (22).

Here we describe studies aimed at assessing the early steps of unfolding of BVPrP^C^E200K (90-231) using the same thermal unfolding experimental design and the same analytical approach that we used for the wt BVPrP^C^(90-231) general case (12).

Our experiments show that the early steps of unfolding are very similar. However, the unfolding pathways diverge somewhat beyond certain temperature. These divergences enrich our understanding of the general process of PrP^Sc^ propagation and also underscore the radical difference between spontaneous, unseeded conversion of PrP^C^ to PrP^Sc^, and PrP^Sc^-templated conversion of PrP^C^ to PrP^Sc^.

## Results

### Thermal unfolding of BVPrP^C^E200K(90-231) shows a lower degree of irreversibility as compared with wt BVPrP^C^(90-231)

We analyzed the thermal unfolding of recBVPrP^C^E200K(90–231) (sequence shown in Suppl. Fig. S1) by monitoring the reduction in the α-helix-associated 222 nm minimum in the far-UV CD spectrum as temperature increased. As shown in Suppl. Fig. S2, the protein unfolded with a melting temperature (Tm) of 61,3 ± 0,13 °C. This is ∼4 °C lower than that of wt recBVPrP^C^(90–231), assessed in parallel for comparison 65,6 ± 1,64 °C and in good agreement with a previous estimate (12).

To assess reversibility, a sample was heated to 75 °C for 1 hour and then cooled to 25 °C. The resulting CD spectrum (Suppl Fig. S2C) showed changes with respect to the initial spectrum acquired at 25 °C with the fresh sample. Specifically, there was a blunting of both the 222 and 208 nm minima, with a new minimum emerging at ∼217 nm. These changes, suggestive of a partial loss of α-helical content and an increase in β sheet secondary structure are similar but substantially milder than those observed when wt recBVPrP^C^(90–231) was subjected to a similar treatment. In that instance the post-75 °C CD spectrum of the wt exhibited a more marked decrease in overall intensity (12) not seen with the E200K mutant.

We then employed solution NMR spectroscopy to obtain detailed, residue-specific insights into the thermal unfolding fingerprint. First, a heteronuclear single quantum coherence (HN-HSQC) spectrum of a sample of freshly folded ^13^C,^15^N-labeled recBVPrP^C^(90–231) dissolved in 10 mM sodium acetate buffer at pH 5, was recorded at 25 °C. Peak assignment was performed by comparison with data from wtBVPrP^C^(90-231) (12). Very minor spectral differences were observed. To resolve ambiguities arising from these differences, we conducted triple resonance experiments, enabling unambiguous assignment of backbone uncertain signals (Suppl. Fig. S3A).

We subsequently recorded a series of N-HSQC spectra at temperatures ranging from 25 °C to 75 °C to investigate temperature-dependent structural changes in recBVPrP^C^E200K(90-231). Up to 45 °C, the spectra retained their high signal intensities, narrow peak widths, and well-dispersed chemical shifts. Peak positions shifted gradually (Fig. 1 and Suppl. Fig. S4), which is consistent with the known temperature dependence of protein amide proton chemical shifts (23).

**Fig. 1.**
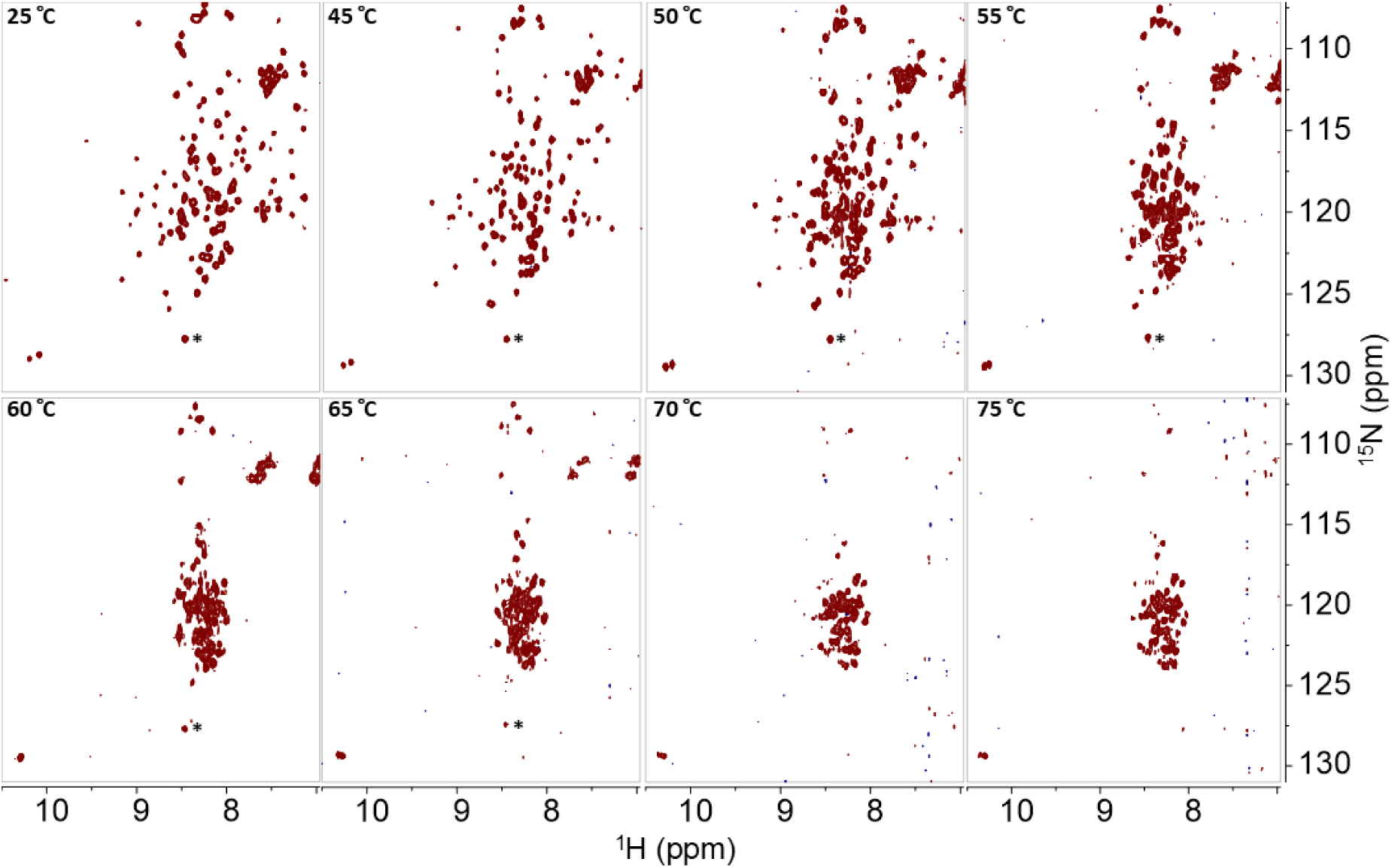
Temperature-induced unfolding of recBVPrP^C^E200K(90-231) tracked with solution NMR. N-HSQC spectra of the backbone amide region from 25 to 75 °C. Each spectrum was recorded at the indicated temperature. The chemical shifts were referenced to the signals marked with the asterisk to facilitate comparison. The spectra are represented at the same noise level.

At ∼50 °C, peaks began to broaden, and several signals diminished in intensity or disappeared entirely (Fig. 1). These alterations became more pronounced with further temperature increases, up to 70 °C. Beyond this temperature, up to the highest temperature tested (75 °C), only minor changes, if any, occurred. These spectra showed broadened peaks grouped in the δ_H_ ∼8.00-8.75 ppm region, characteristic of unfolded and/or disordered secondary structure.

To evaluate the reversibility of these changes, after being heated at each temperature for one hour, the sample was cooled back to 25 °C and another HSQC spectrum was recorded. Similar to the wt protein (12), the temperature-induced changes showed reversibility in the range 25 to 45 °C. However, spectral changes and irreversibility were observed throughout the 50-75 °C range for the recBVPrPE200K(90-231) protein. In this range, substantial peak broadening and/or a marked decrease in peak intensity was observed. The 25post75 spectrum shows that many of the initial peaks in the natively folded sample lost more than half of their intensity after cooling (Fig. 2). The decrease in intensity was not uniform, rather, it affected some signals more than others, and some disappeared completely (Suppl. Fig. S3). This behavior is similar to that of the wt protein, that exhibited irreversible changes beginning at ∼50 °C that were interpreted as the formation of oligomers of ∼12 PrP units (12). Nevertheless, for E200K the loss of peak intensity is substantially smaller, which may indicate a lower putative conversion rate into oligomers (see below evidence of their generation) compared with the wt protein when the sample is subjected to the same heating–cooling cycles.

**Fig. 2.**
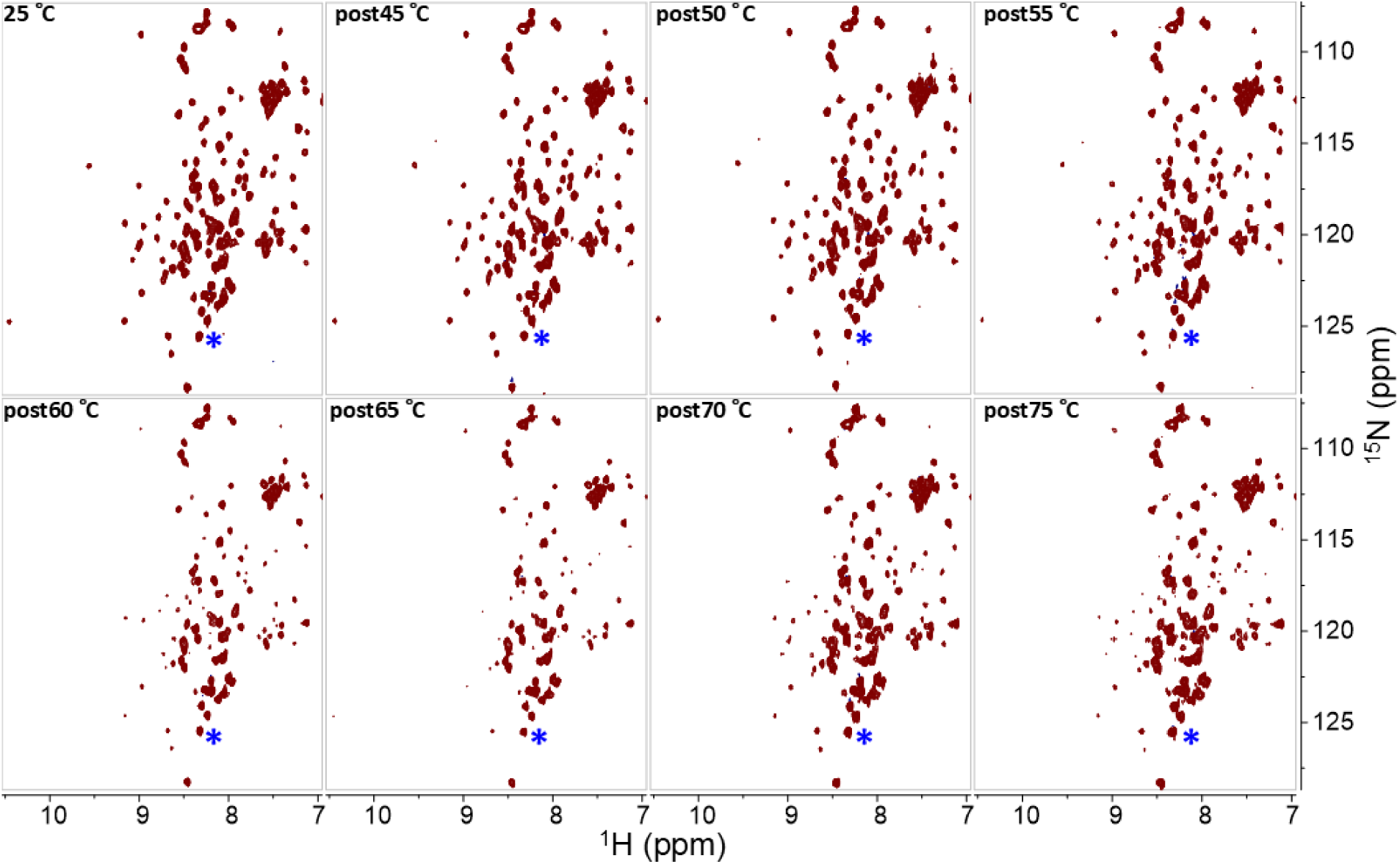
Reversibility of changes in ^15^N-HSQC spectrum of BVPrP^C^E200K(90-231) upon heating. Spectra recorded after heating the sample at the indicated temperatures for 1 hour, followed by cooling at 25 °C. The spectra are represented at the same intensity respect to the signal labelled with the asterisk corresponding to residue V122.

### Temperature dependence of amide H-N chemical shifts identifies likely early BVPrP^C^E200K(90-231) unfolding events

Given that recBVPrP^C^E200K (90-231) remains structurally reversible up to 45 °C under our experimental conditions, we examined residue-level dynamic changes within the 15-45 °C temperature range, using the same experimental approach that we previously developed to probe dynamic changes in wt recBVPrP^C^(90-231) (12). Specifically, we used three solution NMR methods targeting backbone amide HN signals. Each of these methods offers complementary information reporting on the “mobility” (used here in a broad sense) of individual residues, *i.e*., motions in different time regimes, spanning from picosecond to millisecond timescales. Together, they allowed us to identify the most “mobile” regions within the FD, and therefore to build a tentative mapping of regions that may be predisposed to undergo early partial unfolding. More precisely, we analyzed ^15^N relaxation parameters (heteronuclear NOE and R_2_/R_1_ relaxation rates), amide temperature coefficients (Tc), and linearity/non-linearity of Combined Chemical Shift Differences (CCSD) of amide HN groups.

Relaxation in NMR refers to the processes by which nuclear spins return to their equilibrium state after having been excited by dissipating energy to their surroundings. Relaxation rates are sensitive to molecular motions, with fast, small-amplitude motions typically slowing relaxation, whereas slow motions, as in rigid environments, enhance it. We quantified ^15^N longitudinal (R_1_) and transverse (R_2_) relaxation rates and steady-state ^15^N{^1^H} NOE values. These observables are sensitive reporters of motions experienced by backbone NH groups in the ps to ms and ps to ns timescales in solution (24). Residues with low R_2_/R_1_ ratios or low ^15^N{^1^H} NOE values are identified as highly flexible or dynamically disordered, while larger values suggest relative rigidity with a higher contribution of slower motions (12, 25, 26).

As seen in Fig. 3A, B, the same two clear segments observed in similar studies of wt recBVPrP^C^(90-231) can be distinguished in recBVPrP^C^E200K(90-231) according to their ^15^N NOE outputs at the two temperatures tested (25 and 45 °C): the ∼90-120 segment displays negative values with the exception of a stretch of 10 residues with positive values at its center (∼Q98-N108). In contrast, the ∼121-231 segment is characterized by positive values with a few exceptions. Like in wt recBVPrP^C^(90-231), the boundary between these two disordered/ordered subdomains is characterized by a group of residues (∼S120-Y128) with values that cross the zero line and slowly and steadily increase until reaching positive values of the same magnitude as the mean value of FD residues (12). These residues are therefore neither fully ordered nor disordered, showing increasing rigidity from the N to C ends of the ∼A120-Y128 segment.

**Fig. 3.**
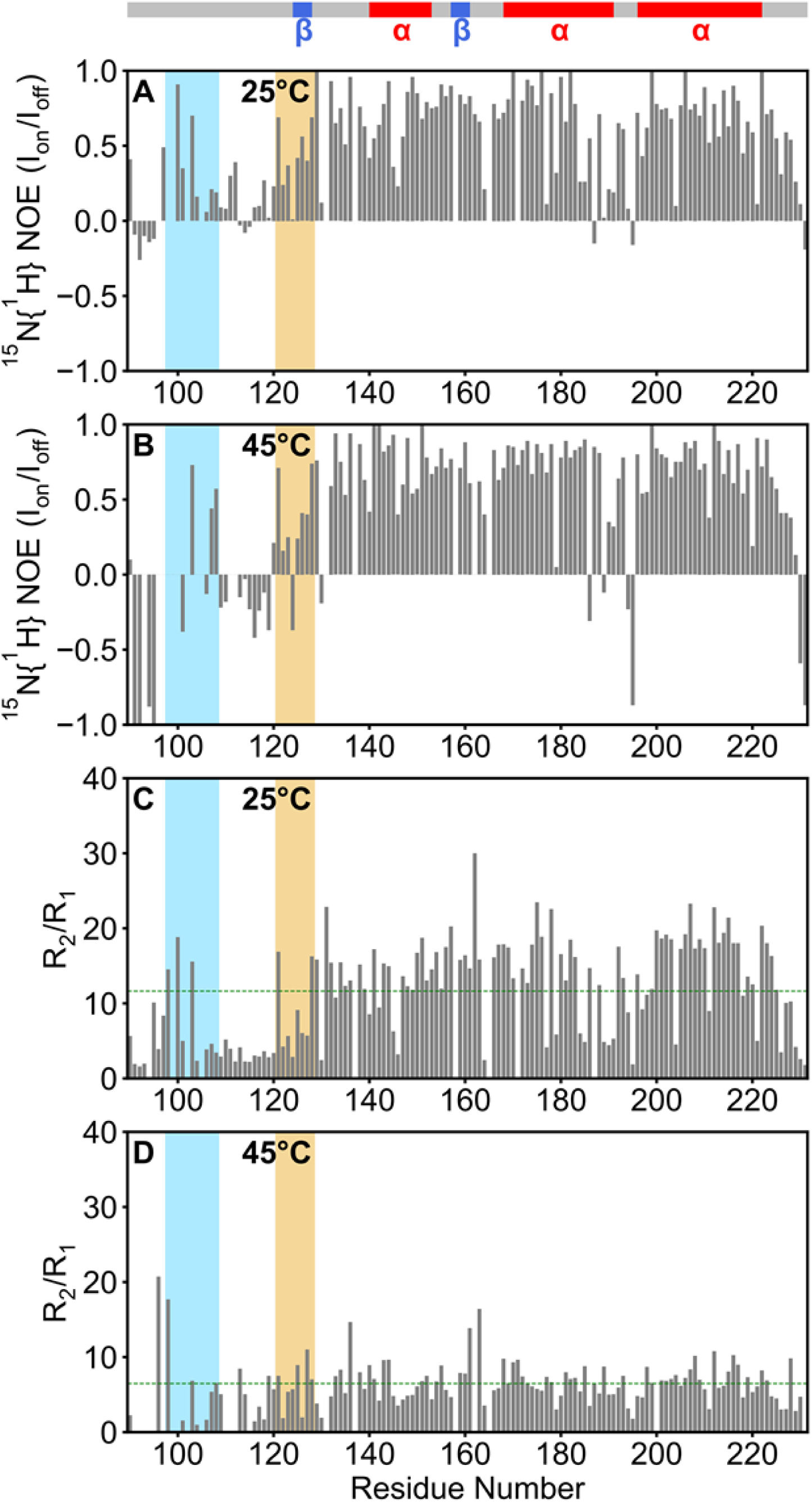
^15^N-NMR relaxation analysis of BVPrP^C^E200K(90-231). ^15^N NOE at **A:** 25 °C and **B:** 45 °C; R_2_/R_1_ ratios at **C:** 25 °C and **D:** 45 °C. A transition zone with increasing ^15^N NOEs and R_2_/R_1_ values, respectively, corresponding to the ∼121-128 stretch is conspicuous (orange rectangles). This sequence is located at the border between the disordered 90-120 and fully folded ∼129-231 domains (see main text). Blue rectangles highlight a stretch of residues within the disordered tail that exhibit unusual ^15^N NOEs and R_2_/R_1_ values, suggestive of transient local order.

R_2_/R_1_ plots at 25 or 45 °C show also conspicuous signatures of the two subdomains according to their group dynamics, faster and slower mobilities. Residues in 90-120 exhibit very low R_2_/R_1_ values (<5), indicative of fast internal motions, whereas residues within the ∼128-225 segment have values in the 10-20 range, typical of well folded domains. As seen for ^15^N NOE, a transition zone with increasing values is seen for the residues at the boundary between these two segments (∼V122-Y128) (Fig. 2C, D).

Relaxation data of residue G131, located at the end of β1, could not be computed, except for R_2_/R_1_ at 25 °C, which was high (Fig. 2). This was so because G131 is affected by intermediate conformational exchange, which substantially elevates its R₂ and causes severe line broadening, reducing its intensity nearly to the noise level (see Suppl. Fig. S2), which is in itself a signal of high mobility in the microsecond to millisecond regime. As a consequence, most of its relaxation data could not be determined. At 45 °C, V161 and Y163 located in β2 also exhibit the markedly high R_2_/R_1_ ratios.

The backbone amide HN protons can exchange with the water protons. This results in a chemical shift drift in the HN signal in the direction of the chemical shift of the H_2_O peak, *i.e*. towards lower ppm. Such drift varies linearly with the rate of this exchange process. Tc values represent the magnitude of the drift as a function of temperature.

Generally, less stable hydrogen bonding associates with backbone mobility (12, 23, 27, 28). We calculated the Tc values of backbone amide proton signals with linear behavior in the N-HSQC spectra over the 15-45 °C temperature range (*vide infra*). As shown in Fig. 4, the Tc plot of recBVPrP^C^E200K(90-231) is essentially similar to that obtained for wt recBVPrP^C^(90-231). As expected, most residues within the disordered N-terminal tail (residues 90-120) exhibit Tc values more negative than -4.6 ppb/K. This implies the absence of stable intramolecular hydrogen bonding consistent with its high dynamic behavior and structural disorder seen by MD simulations (12, 23, 27, 28 and references therein). As with the disordered tail of wt recBVPrP^C^(90-231), the exception is the ∼H96-N100 segment, which likely adopts some transient structure. Residues H96, I139, N143, I182 and M213 exhibit positive Tc values, which is suggestive of temperature-induced conformational change (28, 29).

**Fig. 4.**
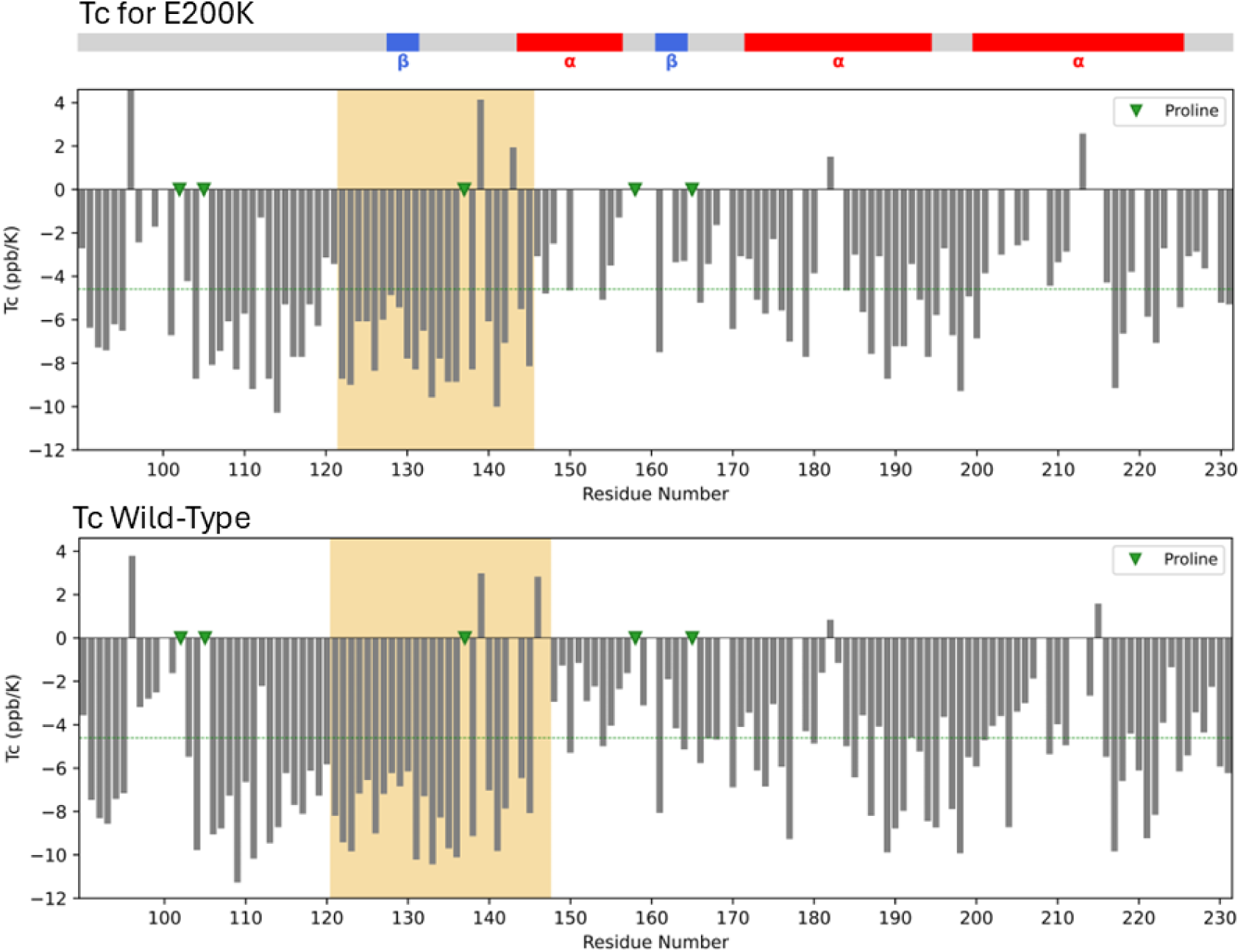
Temperature coefficients (Tc) of backbone amide protons of BVPrP^C^E200K(90-231). The bar plot shows the temperature coefficients T_C_ (ppb/K) of amide proton (HN) signals in the 15-45 °C range represented along the primary sequence. The Tc values of residues giving bad linear fits have been omitted. Signals with values Tc > –4.6 ppb/K are considered protected from the solvent or involved in permanent hydrogen bond and vice versa. **Top:** plot of BVPrP^C^E200K(90-231) data. **Bottom:** plot of wtBVPrP^C^(90-231) data reproduced for comparison from Sabzehei et al. (12) with permission from the authors. The region highlighted in orange in both plots represents a continuous stretch of residues with Tc < –4.6 ppb/K values contiguous to the flexible 90-120 tail (see main text). It is slightly shorter in BVPrP^C^E200K (90-231). The secondary structure elements are shown in the top bar; green triangles indicate proline residues, for which Tc cannot be determined because they lack an amide HN.

Residues from V122 to W145, which include the V122-G127 transition coil (as identified with ^15^N NOE and R_2_/R_1_ measurements), β1, the coil connecting β1 and α1, and the N-terminal edge of α1, form an uninterrupted segment, ∼V121-E144, with strongly negative (or positive) Tc values (Fig. 4), consistent with weak intramolecular hydrogen bonds, interpretable as a propensity to unfold. Note that this segment is just slightly shorter than the equivalent high unfolding propensity V121-E146 segment in wt recBVPrP^C^(90-231) (Fig. 4). Also to note in the mutant protein residues A120 and V121 exhibit Tc values indicative of hydrogen bonding and/or relative rigidity, in contrast with their counterparts in the wt protein, and therefore in the mt protein the A113-D144 continuous highly negative segment is interrupted.

As in the wt protein, the remainder of α1 (essentially all of it in this case), and the short coil connecting it to β2, display Tc values less negative than -4.6 ppb/K, indicating greater stability, whereas residues in β2 and in the loop connecting β2 to α2, show a mix of high and low Tc values, suggesting structural complexity.

Also, as in the wt protein, both α2 and α3 in recBVPrP^C^E200K(90-231) display a pattern of Tc values less negative than -4.6 ppb/K in their central regions (*i.e*., relative rigidity), around the disulfide bond connecting the two helices, and more negative toward their N- and C-termini, indicating greater relative flexibility of these regions, and the G195-T199 coil connecting α2 and α3 exhibits highly negative Tc values, suggesting disorder.

Furthermore, as with wt recBVPrP^C^(90-231), we analyzed the temperature-dependent (VT) changes in amide ^1^H and ^15^N chemical shifts using N-HSQC spectra (Suppl. Figs. S4, S5). As a protein is heated, the resulting increased thermal motion typically accelerates conformational averaging, and the protein is more amenable to overcome conformational energy barriers. Heating also increases the exchange rate of amide protons with water protons, leading to upfield shifts in both ^1^H and ^15^N chemical shifts (23) that are more pronounced for residues not involved in intramolecular hydrogen bonds. These changes are typically linear with respect to temperature. However, deviations from linearity are associated to residues that sample alternative, higher-energy, and less populated conformations, *i.e.,* a form of mobility (27, 28, 30).

Thus, following the approach we previously used to probe wt recBVPrP^C^(90-231), we combined for each peak the ^1^H and ^15^N chemical shift drift at a given temperature into a single value called the Combined Chemical Shift Difference (CCSD; Eq. 2) across the 15-45 °C range. The temperature dependence of each CCSD was evaluated by fitting the data to a linear regression model (Eq. 3). This assessment of linearity is relevant for our objectives as it correlates with the conformational stability of the residue (12, 31, 32). Residuals, representing deviations from linearity, were then plotted against temperature (Suppl. Fig. S4).

Residues showing clear non-linear behavior were identified by visual inspection of these plots, and quantitatively by linearization of the residuals and calculation of their curvature.

In contrast with the Tc results, that were very similar between recBVPrP^C^E200K(90-231) and wt recPrP^C^(90-231), some differences were seen in CCSD values corresponding to these two PrP sequential variants. Most residues in the FD showed a linear dependence of CCSD with temperature, which indicates conformational stability. Significant deviations (non-linear CCSD residuals) were observed in a few mostly isolated residues, such as M129, located within β1 (but not in any of the other three residues of β1). None of the four residues in β2 was non-linear either. Other non-linear residues were: N143, Q160, Y169, H177, T184, Q186, T193, K204 and E221. This contrasts with the larger number of non-linear residues in the wt recPrP^C^(90-231): V121, V122, L125, H140, N159, C179, I184, T193, K194, G195, T199, D202, V203, V210 M213 and Y226 in the wt recPrP^C^(90-231). It thus appears that while more or less the same regions contain residues with non-linear CCSD values, fewer residues (9 *vs*. 16) and in cases not exactly the same residues have non-linear values in recBVPrP^C^E200K(90-231) (Suppl. Fig. S5).

No residues in the very high mobility 90-120 disordered tail feature non-linear CCSD plot. This can be ascribed to the fact that residues that explore a large number of conformational states with rapid interconversion typically display a linear CCSD-temperature relationship, as solvent interactions dominate (33). This trend is also apparent in the disordered 227-231 C-terminal tail, where no non-linear residues were detected. Similar results were obtained for the wt protein (12).

It should be noted that, within the temperature range used herein for Tc and CCSD studies reported, all backbone amide signals were in fast conformational exchange in the chemical shift timescale, with the exception of residues G131, V161 and Y163, that showed intermediate exchange at some temperatures.

### At high temperatures, BVPrP^C^E200K(90-231) unfolding intermediates behave differently from those of wtBVPrP^C^(90-231)

As mentioned above, many signals in the N-HSQC spectrum began to broaden beyond approximately 50 °C (Fig. 1), becoming less intense and eventually completely vanishing. At 75 °C, the spectrum contained only a fraction of the peaks observed in the 15-45 °C temperature range.

However, in contrast with the behavior of wt recBVPrP^C^(90-231), when the samples were cooled back to 25 °C, many of these changes did not persist, and the recovered spectrum looked relatively similar to the original one, indicating that these changes were reversible (Fig. 2). To further investigate this, we performed size exclusion chromatography (SEC) on a post-75 °C sample. A freshly prepared wt recBVPrP^C^(90-231) post-75 °C sample was analyzed in parallel using the same column and settings, for comparison. The chromatogram of the mutant protein shows, together with the peak corresponding to the recBVPrP^C^E200K(90-231) monomer, a peak with a shorter elution time similar to that of the oligomeric species previously described for wt recBVPrP^C^(90-231) (Fig. 5A), that based on calibration of the column corresponds to a ∼12mer. However, the area of this peak was substantially smaller than that of the wt protein: it corresponded to 14% of the total area, as compared with 41% in the wt sample (Suppl. Fig. S6). In addition, another small peak eluting very near the void volume, and therefore corresponding to a much larger oligomer and representing 9% of the total area was seen (Fig. 5A and Suppl. Fig. S6). Such peak was absent in the post-70 °C wt sample, as previously reported (12). In agreement with these results, negative stain Transmission Electron Microscopy (TEM) of the post-75 °C mt sample revealed spherical particles with a very similar appearance to those seen in the wt sample (Fig. 5B,C,D) but in a much lower abundance.

**Fig. 5.**
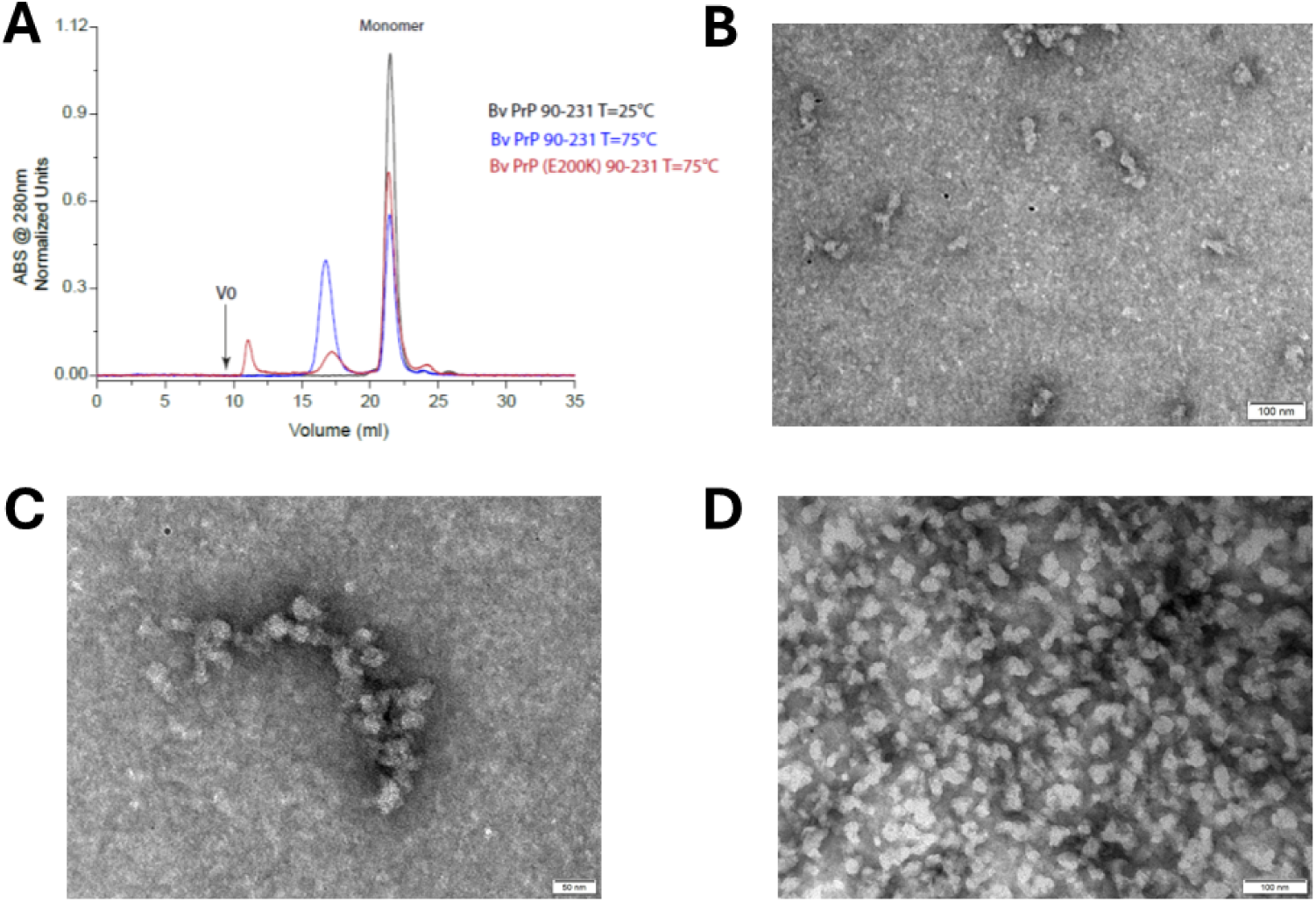
Partially unfolded recBVPrP^C^E200K(90-231) undergoes oligomerization dynamics above 50 °C that differ from those of the wt protein. **A:** SEC analysis of recBVPrP^C^E200K(90-231) and wt recBVPrP^C^(90-231); V0 = void volume. **B:** Negative-stain TEM image of recBVPrP^C^E200K(90-231) after heating to 75 °C for 1 h followed by cooling to RT. **C:** Same as B, shown at higher magnification. **D:** Negative stain TEM image of wt recBVPrP^C^(90-231).

Having established the presence of oligomers, both ∼12-mers and larger ones, we considered interrogating, at the residue level, the changes seen in the HSQC spectrum of the post-75 °C sample (Fig. 2 and Suppl. Fig. S3), by calculating post75 °C/25 °C signal intensity ratios as we did for the wt sample (12). However, the fact that the predominant species in the post75 °C (≥72%) were monomers, dissuaded us, as results would hardly be representative of the contribution of oligomers.

### Spontaneous conversion of recBVPrP^C^E200K to PrP^res^ *in vitro* using Protein Misfolding Shaking Amplification (PMSA)

**Rec**BVPrP^C^E200K(90-231) and wt recBVPrP(90-231) were subjected to PMSA in the absence of seed using glass beads, known to facilitate spontaneous conversion of recombinant PrP substrates to a PrP^res^ conformation using dextran sulfate as the sole cofactor (34, 35). It is noteworthy that recBVPrP^C^(23-231) is one of the most readily spontaneously converted recombinant PrP variants among the hundreds tested, which together represent all known PrP variants described across mammalian species (36). Wt recBVPrP^C^(90-231) displayed a spontaneous misfolding proneness that was considerably lower than the one reported for the full-length protein: only 2 out of 8 replicates developed PK-resistant PrP (PrP^res^) in the first passage and 3 out of 8 in the second (Fig. 6), compared with 8/8 and 8/8 for the full-length species (36). Remarkably, recBVPrPE200K(90-231) converted spontaneously in the same proportion of vials, 2/8 and 3/8 respectively (Fig. 6) indicating that the mutation does not confer a higher propensity for spontaneous misfolding. The experiment was repeated under identical conditions with equivalent results, confirming that the spontaneous misfolding capacity of both proteins is comparable.

**Fig. 6:**
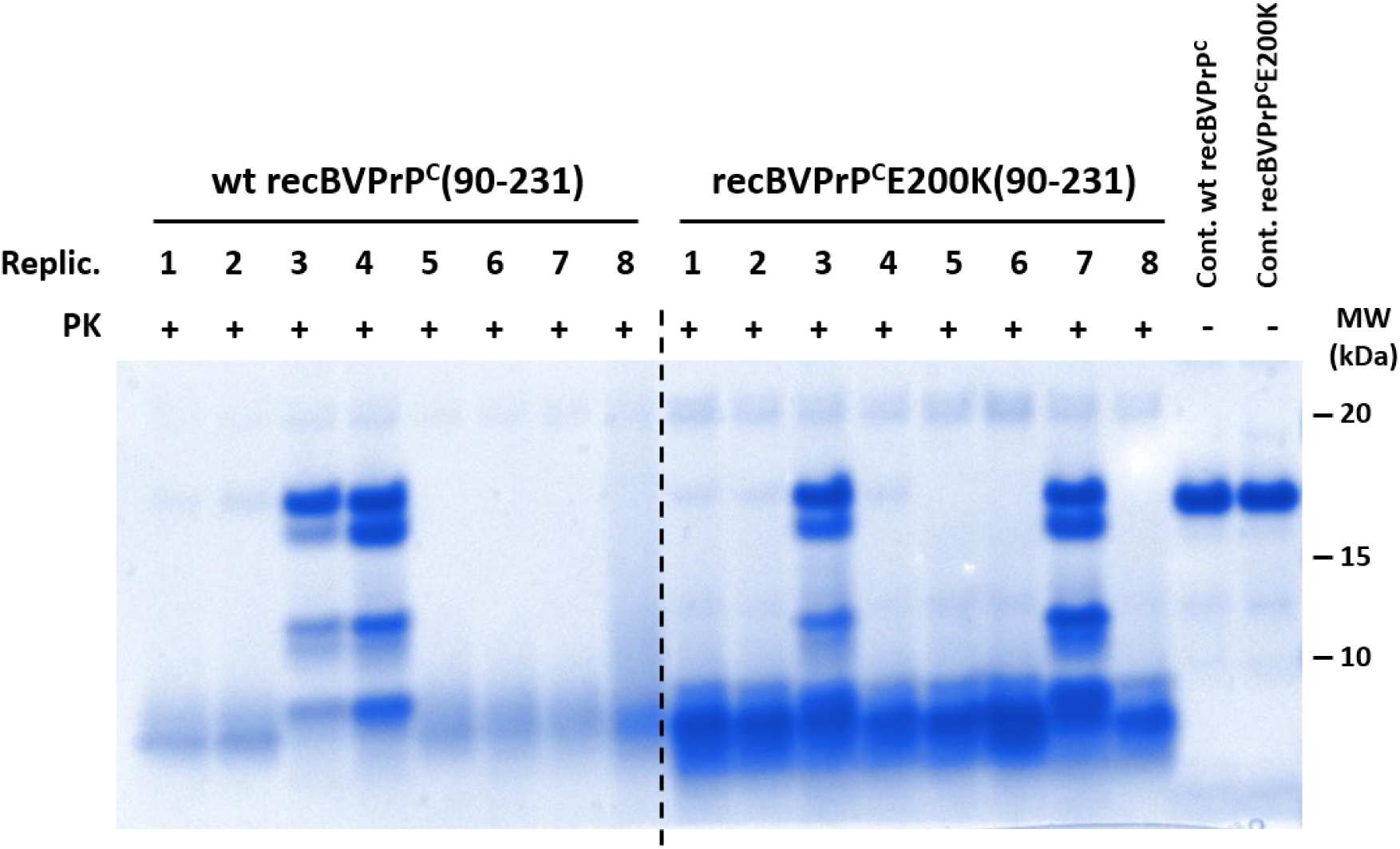
Rec BVPrP^C^200K(90-231) does not have a higher propensity to spontaneously convert to PrP^res^ in vitro through PMSA than wtBVPrP^C^(90-231). wt recBVPrP^C^(90-231) and recBVPrP^C^200K (90-231) were subjected to PMSA with no seed in the presence of 1 mm diameter (replicates 1-4 of each variant) and 0.1 mm diameter (replicates 5-8 of each variant), i.e. 4 replicates for each condition, 8 per protein variant. Following PMSA (see ref. 36), the products were treated with PK, subjected to SDS-PAGE and gels stained with Coomassie blue.

The lower conversion efficiency of both truncated variants relative to full-length wt recBVPrP(23–231) may not be attributable solely to the absence of the N-terminal region (23–89), which contains the octapeptide repeat and polybasic segments that favour the interaction with polyanionic cofactors. The His-tag retained in these constructs might also contribute, as it may sterically hinder the conformational rearrangements required for conversion and/or modify the electrostatic interaction with dextran sulfate, thereby reducing the efficiency of spontaneous misfolding.

## Discussion

We have recently proposed that, given the dull nature of PrP^Sc^, the FD of a converting PrP^C^ unit must unfold before it can be templated by PrP^Sc^ (12). This proposal is based on the view that PrP^Sc^ adopts a classic, essentially rigid PIRIBS amyloid architecture (4-8), whose templating surface undergoes only random motions (37) and whose apparent templating capacity, as in other classic amyloids, is limited to the docking and templating of previously unfolded protein sequences.

Partial unfolding of the PrP^Sc^-tethered FD could follow as a consequence of intrinsic “breathing” motions of FD and/or be facilitated by physical properties of the FD local micro-environment, interactants or cellular cofactors. But in any case, the PrP^Sc^ templating surface would participate in a limited way, acting as a mere “Procrustean bed” onto which unfolded segments of the FD can be trapped and converted (12). We have assessed thermal unfolding of BVPrP^C^(90-231), a sequence chosen because BVPrP^C^ can adapt to all known PrP^Sc^ strain conformations (38, 39), *i.e*., it is a universal converter to PrP^Sc^. Our data strongly suggested that for the wt protein the ∼121-144 stretch, *i.e*., the region of the FD immediately contiguous to the disordered ∼90-120 domain, that is the first to convert, is the most likely one to unfold and continue the conversion process, both for reasons of intrinsic plasticity and accessibility of and proximity to the templating surface (12). Here we have mapped early unfolding events of the E200K mutant using thermal unfolding and identified the ∼121-144 segment also as the most likely region to unfold first in recBVPrP^C^E200K(90-231), just as is the case in wt recBVPrP^C^(90-231) (12). This conclusion is based on 1) the Tc of backbone amide protons, that largely reflects the stability of the hydrogen bond it forms. As seen in Fig. 4A, the most prominent continuous stretch of residues with Tc < –4.6 ppb/K or Tc > 0 ppb/K spans V122 to W145 (Fig. 4). This is equivalent to the V121-E146 stretch in wtBVPrP^C^(90-231). Furthermore, the next contender for high mobility, the segment around the α2-α3 loop, contains fewer residues with Tc < –4.6 ppb/K as compared to wtBVPrP^C^(90-231) (Fig. 4). It is noteworthy that V121 does not exhibit a Tc < –4.6 ppb/K value in the mutant protein, as it does in the wt one. 2) Relaxation data are comparable with those obtained for the wt, with some nuances. In both proteins there are the expected two clear segments, connected by a transition stretch of residues. The ∼122-126 stretch displays intermediate values of ^15^N(^1^H) NOE and R_2_/R_1_ in both the mt and wt proteins, which can be interpreted as evidence of very high flexibility not reaching the characteristics of a fully disordered state. Beyond this stretch, β1 (Y128-G131) also appears to be particularly labile. G131 was barely visible in the HSQC spectrum at 25 °C, (Suppl. Fig. S3), suggesting high mobility in the microsecond to millisecond regime. This in turn led to unreliable ^15^N(^1^H) NOE values, both at 25 and 45 °C, and a high R_2_/R_1_ ratio at 25 °C (Fig. 3). Also, M129 exhibited a non-linear CCSD (Suppl. Fig. S5). In this respect, it is noteworthy that non-linearity of CCSD temperature dependence of residues within BVPrP^C^E200K(90-231) differs somewhat with that of residues within wtBVPrP^C^(90-231). Deviations from linearity of the CCSD are interpreted as exploration of alternative conformations in the fast-exchange regime by the residue involved, *i.e.,* mobility (12, 31, 32). While there was a considerable coincidence of residues with non-linear CCSD temperature dependence, the number of non-linear residues in the mutant protein was only ∼half of the number in the wt protein (12 and Suppl. Fig. S5). Very relevantly, while in the wt, 3 of 4 residues in β1, and 2 of 4 in β2, are non-linear, in the mt only 1 residue in β1 (M129) and none in β2 were (Suppl. Fig. S5). This might suggest that disruption of the β1/β2 assembly perhaps are not so prominent as a first event as in the wt.

These conclusions are derived from data obtained in the 15-45 °C temperature range, in which only “breathing”, or immediately reversible motions take place, as surmised from intact dispersion and intensity of peaks in the H-N HSQC spectra (Fig. 1). These changes are fully reversible, as seen by full recovery of the 25 °C spectrum upon cooling to this temperature. In the wt protein, 45 °C marks the limit of reversibility, and beyond this temperature, more clearly beyond 50 °C (there is a grey transition zone between 45 and 50 °C), changes become irreversible as a consequence of progressive oligomerization (12). In the mt protein, reversibility was more resilient, with much more modest permanent peak broadening and intensity loss seen upon cooling after heating at 75 °C (Fig. 2), and less pronounced permanent changes in the CD spectrum after cooling (Suppl. Fig. S2). It should be noted that even more marked reversibility of temperature-induced unfolding has been described for other PrP proteins, in particular HuPrP^C^(90-230), as reported by Russo *et al*. These authors observed that HuPrP^C^(90-230) undergoes complete unfolding in a virtually fully reversible way (40).

In our earlier publication we interpreted the soluble β-sheet-rich oligomers formed above 50 °C by wtBVPrP^C^(90-231) as useful artifacts capturing a transient late stage unfolding intermediate that exists just instants before collapsing into the oligomeric state and hypothesized that in the presence of the PrP^Sc^ template, such unfolded conformers, tethered as they would be to PrP^Sc^ through their N-termini, would collapse onto the templating surface, rather than onto similar partially unfolded PrP^C^ companions. While the majority of signals in the H-N HSQC spectrum arising from the FD disappeared in the oligomer-rich samples, a substantial number of signals from α2 and some from α3 remained, supporting the notion that the β2-α2-α3 segment of the wtBVPrP^C^(90-231) is likely the last portion of its FD to unfold and convert (12). The less pronounced reversibility of thermally induced changes above 50 °C in the mt protein is likely due to the much lower amount of soluble oligomers formed (Fig. 5 and Suppl. Fig. S6).

Under conditions similar to those used in our experiments, ovine PrP(23-231) has been shown to oligomerize like BVPrP(90-231) during thermal unfolding (41-43), whereas HuPrP(90-231) and HuPrP(23-231) did only when heated at 61 °C for several hours, and produced only minute amounts of oligomers (40). While the reasons resulting in partially unfolded E200K mt monomers to associate forming oligomers would be interesting to know, this lies outside of the scope of this study.

The E200K mutation does not result in any substantial changes in the conformation of the recHuPrP FD (15). The only relevant change is the absence in E200K mt of the E200-K204 salt bridge at the N-terminal region of α3 that exists in the wt protein (15). However, the mutation does have a deep impact on the distribution of surface charges, particularly at slightly acidic pH. This results in a dramatically altered electrostatic potential in the mutant protein (15). Namely, the E200K mutation breaks the relatively uniform distribution of surface charges, introducing patches of positive potential. Under mildly acidic conditions, the surface of the mutant protein contains separate patches of negative and positive electrostatic potential, whereas the wt protein has a homogeneous, mixed distribution of charges (15). Zhang *et al*. argued that these surface-restricted changes are unlikely to affect conformational transitions in isolated PrP molecules. In contrast, they could have profound effects on the ability of PrP^C^(E200K) to interact with other molecules acting as facilitators of the conversion reaction (15). An increase in the surface negative charge as afforded by the E200K mutation might facilitate binding of anionic cofactors to PrP, and in turn facilitate its spontaneous conversion to PrP^Sc^. In this respect, Marrone and cols. using molecular dynamics (MD) approaches, have proposed a tendency of the E200K mutant PrP to form dimers and eventually polymers, as a consequence of its peculiar surface charge distribution (44). It is tempting to speculate that, if as we have proposed, spontaneous generation of PrP^Sc^ is initiated by anionic cofactor-promoted association and conversion to PIRIBS of lysine-rich disordered ∼90-120 segments (35), the “hanging” FDs might interact more easily if their surface charges are grouped in positive and negative patches, as in the E200K mutant (44), rather than be repelled from one another electrostatically. In any case, none of these considerations applied to our *in vitro* experiment, which showed a similar capacity of the E200K and wt proteins to spontaneously generate PK-resistant PrP under our experimental (PMSA) conditions.

Another structural feature that may be particularly relevant to the E200K substitution is the specific position of E200 at the N terminal end of α3. The α helices possess an intrinsic macrodipole arising from the cumulative alignment of peptide-bond dipoles, resulting in a partial positive charge at the N-terminal end and a partial negative charge at the C-terminal end (45). Consequently, negatively charged residues located near the N-terminus of an a-helix can provide favorable electrostatic stabilization of the helical structure (46, 47). The presence of E200 at this position in wtPrP may therefore contribute disproportionately to the local stability of H3, beyond what would be expected from the intrinsic properties of the residue alone. Substitution of Glu by Lys in E200K not only removes this potentially favorable interaction but introduces a positively charged side chain in a region of positive helix-dipole potential, potentially generating an unfavorable electrostatic contribution. This positional effect was not explicitly considered in the original structural interpretation of E200K (15) and could provide an additional mechanistic explanation for the experimentally observed destabilization of α3. In particular, the loss of this favorable electrostatic contribution may facilitate local unfolding of α3 and thereby significantly lower the kinetic barrier separating the native PrP conformation from conformational states competent for spontaneous conversion. Thus, despite having only a relatively modest effect on the overall thermodynamic stability and native fold of PrP, the E200K substitution could exert a disproportionately large effect on the kinetics of the initial conformational transitions leading to spontaneous PrP conversion.

To sum up, our data show that the BVPrP^C^E200K(90-231) FD exhibits an early unfolding pathway essentially similar to that of wt BVPrP^C^(90-231), with some nuances, in particular a less unstable β1/β2. Once its ∼121-140 segment has unfolded and converted to the PrP^Sc^ conformation, our data do not allow us to predict what might happen. However, given the similar conformation and Tm of the mt and wt PrP proteins, it is tempting to speculate that their ∼137-231 “open” forms (48) might behave similarly. Studies to compare the unfolding of wt and E200K ∼137-231 are in course. Moreover, whether the ∼121-140 segment unfolds in a sequential, zipper-like fashion or not, both in the mt and wt proteins, is currently unknown.

Our data also show that the E200K mutant, despite having a slightly lower Tm (∼4 °C lower) than that of the wt protein, seems to exhibit, paradoxically, a somewhat lower overall flexibility. In particular β1/β2 but also the α2-α3 connecting loop and adjacent segments exhibit overall, less negative T_C_ values. Therefore, it might be the case that, counterintuitively, the E200K mutant be a less effective propagator of PrP^Sc^ than its wt counterpart. As mentioned earlier, such comparison would in any case be very difficult to establish, given that to eliminate propagation barriers the meaningful comparison would be conversion of mt PrP^C^(E200K) in the context of mtPrP^Sc^(E200K) templating with conversion of wtPrP^C^ in the context of wtPrP^Sc^ templating. Such a comparison might be attempted using the 1x BVPrP wt and E200K mice recently described by Watts and cols. (49), but it is outside the scope of the present study. This leaves its higher intrinsic spontaneity (*i.e*., capacity to misfold into a PrP^Sc^ conformation in the absence of a PrP^Sc^ seed) as the reason why the E200K mutant drives genetic prion disease with an extremely high penetrance (20).

## Materials and Methods

### Recombinant protein production

Recombinant bank vole (BV) PrP (90–231), corresponding to the I109 natural polymorphism (GenBank accession number PQ327920), was produced in both wild-type (WT) and E200K forms. The open reading frame of the wild-type I109 bank vole *Prnp* gene was obtained from genomic DNA by PCR and cloned into the pOPIN E expression vector, which adds an amino-terminal 6×His tag to the encoded protein, without any linker. The E200K variant was generated on the same I109 background by GenScript through gene synthesis followed by subcloning into the pOPIN E expression vector using the CloneEZ cloning method. WT and E200K proteins were expressed and purified following the same procedure as described previously (50), with WT used as a control in the comparative experiments.

We decided to keep the N-terminal 6×His tag used to facilitate purification for practical reasons, like others before us (51-55). It has been experimentally established that the 6xHis tag, at the end of the intrinsically unfolded ∼90–120 segment, does not influence unfolding of the FD (52). *E. coli* Rosetta™ (DE3) competent cells were transformed with the expression vector using standard molecular biology procedures. Expression of ^13^C,^15^N labelled recombinant protein was achieved by growing the bacteria in two flasks with 50 ml each of LB broth with 50 μL of ampicillin overnight and then for 4–5 h in two 2 L Erlenmeyer flasks containing each 500 ml of minimal medium supplemented with 3 g/L of uniformly labelled ^13^C-glucose and 1 g/L of ^15^N-NH_4_Cl. Upon isopropyl β-D-1-thiogalactopyranoside (IPTG) induction, inclusion bodies containing the protein were harvested. Purification of the protein was performed with a histidine affinity column (HisTrap^TM^ HP crude 5 ml, Cytiva Sweden AB). After elution in buffer consisting of 20 mM Tris-HCl, 500 mM NaCl, 500 mM imidazole and 2 M guanidine-HCl, pH 8, the quality and purity of protein batches were assessed by BlueSafe (NZYtech) staining after electrophoresis in SDS-PAGE gels (BioRad). Finally, guanidine-HCl was added to a final concentration of 6 M for long-term storage of purified protein preparations at −80 °C.

Protein folding was performed by dialysis against 10 mM sodium acetate, pH 5, using 3 kDa cutoff membrane Slide-A-Lyzer dialysis cassettes (Thermo), with three changes of dialysis buffer. Aggregated material was removed by a brief centrifugation at 18,000 × *g* in a Beckman Microfuge 22R tabletop centrifuge. Protein concentration was estimated spectrophotometrically using an ε_280_ of 26,000 and a MW of 17.02 kDa. The folded protein was concentrated as needed using Amicon centrifugal devices and immediately used for different analyses or kept at 4 °C for a few days between experiments. In such cases, the quality of the protein was assessed by ^1^H–^15^N HSQC spectra.

### Far-UV Circular Dichroism (CD)

For thermal unfolding experiments, CD spectra were acquired in a Jasco 1100 spectrometer with temperature control at a 0.7 mg/ml protein concentration, in 10 mM sodium acetate, pH = 5 placed in a 1 mm light-path quartz cuvette. A temperature ramp from 25 to 85 °C was applied, with spectra between 180 and 260 nm acquired every 1 °C. Five spectra were acquired and accumulated for each data point. For comparative spectra at 25 °C, samples were diluted to a protein concentration of 0.1 mg/ml and CD spectra acquired using a Jasco715 spectrometer and 1 mm light-path quartz cuvettes. Twelve spectra were acquired and accumulated.

CD data as a function of temperature was fitted by SciDavis software to a single-state sigmoidal Boltzmann model to calculate the percentage of unfolding (Eq. 1).

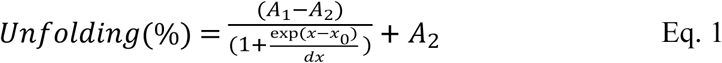

where *A_1_* and *A_2_* are the minimum and maximum value, *x_0_* is the midpoint of the transition and *dx* is the time constant.

### Size Exclusion Chromatography (SEC)

Samples subjected to different temperatures (*vide supra*) and cooled back to 25 °C. After that they were subjected to SEC using a 90 cm TSK4000SW (7 mm×600 mm) column eluted with 10 mM sodium acetate at 1 ml/min. A_280_ of the effluent was monitored. This column was previously calibrated for molecular mass using low and high molecular mass calibration kits (Amersham). Before each run the column was equilibrated with at least four column volumes of elution buffer. Each run was performed at 20 °C (42).

### Transmission electron microscopy

BVPrP90-231 (10 μl) samples were adsorbed on freshly glow discharged 400 mesh carbon-coated copper grids (Electron Microscopy Sciences), washed with water twice, negatively stained with freshly filtered 2% uranyl acetate for 20 s, air-dried and imaged using a JEOL JEM 1011 electron microscope at 100 KV and a MegaView G3 camera.

### NMR spectroscopy

All NMR experiments were carried out in solution using a Bruker NEO 750 MHz spectrometer equipped with a PA-TXI-HFCN (^1^H-^19^F/^13^C/^15^N) room temperature probe with an automated tuning and matching system, and temperature control for VT. The spectrometer control software was TopSpin 4.3.x.

### NMR sequence assignments

The protein, in 10 mM sodium acetate buffer, pH 5, was supplemented with D_2_O (10%). The final protein concentration was 6 mg/ml. The assignment of Ca, Cb, C, N, HN and Ha resonances at 25 °C was carried out with the following 3D standard triple resonance experiments: HNCA, intra-HNCA, HNCACO, CBCANH, intra-CBCANH, CBCA(CO)NH, HNCO, HN(CO)CA, HNHa, HNCAHA, HCCH-TOCSY, ^15^N and ^13^C edited TOCSY-HSQC and ^15^N and ^13^C edited NOESY-HSQC. The spectra were processed with TopSpin and analyzed with CCPN software v3.2 (56) guided by the chemical shifts previously reported for wtBVPrP(90-231) (12) and deposited under the BMRB accession code ID:53716. The chemical shift assignment of N and HN resonances at different temperatures was carried out by tracking peak trajectories in a series of N-HSQC Spectra (*vide infra*) over the temperature range. Spectra were processed with TopSpin and the peak positions were analyzed using MestreNova software (Mestrelab Inc.) ^1^H, ^13^C and ^15^N chemical shifts were referenced indirectly to external 2,2-dimethyl-2-silapentane-5-sulfonic acid (DSS) reference.

### Evaluation of Temperature coefficients (T_C_)

Temperature coefficients were determined by analyzing the chemical shift of the amide proton (28 and refs. therein) of each residue in a series of N-HSQC spectra measured in the temperature range of 15 °C to 45 °C, in 5 °C or 2.5 °C increments. The HN chemical shift versus temperature for each amino acid was linearized by MS-Excel software, and the temperature coefficient (*T_C_*, in ppb/K) was taken from the slope. *T_C_* values were plotted along the PrP^C^ sequence.

### Variable Temperature (VT)-NMR data analysis

N-HSQC spectra were acquired at increasing temperatures from 15°C to 45°C in 5°C or 2.5 °C increments. Each spectrum was referenced using the acetate signal of the buffer as secondary reference, and chemical shift changes for each amide proton-nitrogen (HN) cross-peak were tracked across the temperature range. The ^1^H and ^15^N drift of a peak in the N-HSQC spectrum at a certain temperature of the series was analyzed by the Combined Chemical Shift Difference (CCSD) (57) (Eq. 2):

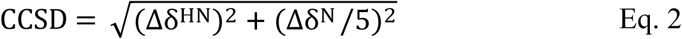

Where Δδ^HN^and Δδ^N^is, respectively, the change in ^1^H and ^15^N chemical shift respect to the reference values of the spectrum acquired at the lowest temperature of the series.

The experimental CCSD for a specific amino acid at each temperature, was linearized using MS-Excel software by fitting the data to the following equation (Eq. 3):

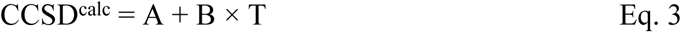

Here, CCSD^calc^ represents the calculated linearized CCSD value for each data point, T is the temperature, and A and B are the regression coefficients. The residual error (defined as CCSD - CCSD^calc^) was calculated and plotted against temperature (31, 32). Residual errors showing linear or non-linear behavior were identified by visually inspecting the plots and quantitatively by curvature calculation through second-order linearization of the residuals and evaluation of the RMSD factor. Plots of the residuals were represented by SciDAVis software (scidavis.sourceforge.net) or by a homemade python script based in the *matplotlib* library. The non-linearities identified in the CCSD study were represented color-coded and overlaid onto the primary structure sequence using Polyview 2D software (58).

### NMR relaxation

^15^N relaxation experiments based on the ^15^N-HSQC detection (25, 59) were carried out at 25 °C and 45 °C at a 6 mg/ml protein concentration. Each spectrum was measured with 8 scans and 128x2048 points in the ^15^N and ^1^H dimensions. The Longitudinal relaxation rates (R_1_) were measured (sequence *hsqct1etf3gptcwg3d*) exploring 12 points of the recovery delay between 2 ms and 1.2 s. Transverse relaxation rates (R_2_) were measured (sequence *hsqct2etf3gptcwg3d*) exploring 12 points of the CPMG duration between 15.6 and 343 ms.

For determining ^15^N *R_1_*and ^15^N *R_2_*, the intensity of a given cross peak in the N-HSQC correlation at each relaxation delay *τ*, *I(τ),* was measured and the curve was non-linearly fit to the mono-exponential equation Eq. 4 to determine *I_0_* and *R_x_*.

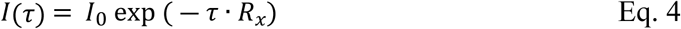

Where *R_x_* refers to either ^15^N *R_1_* or ^15^N *R_2_*, and *I_0_* is the initial intensity at *τ* =0.

Steady state ^15^N{^1^H} heteronuclear NOE (^15^N NOE) (sequence *hsqcnoef3gpsi*) was measured by acquiring two interleaved spectra collected with (*S^sat^*) and without (*S^0^*) an initial proton saturation period (3 s) during the 5 s recycle delay. The ^15^N NOEs were determined by measuring the intensity of a given cross peak in the saturated (*S^sat^*) and reference (*S^0^*) spectra using Eq. 5.

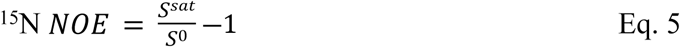

### Spontaneous conversion of PrP^C^ to PrP^Sc^ in vitro using PMSA

Preparation of recombinant PrP based PMSA substrates for spontaneous recPrPres formation, was carried out as previously described (36). Briefly, the purified recPrP stored at −80 °C was diluted 1:5 in phosphate buffered saline (HyClone) and dialyzed for re-folding against PBS at 1:2,000 ratio for 1 h at room temperature. The dialyzed protein was centrifuged at 19,000 × *g* for 15 min at 4 °C to eliminate amorphous aggregates and the supernatant, containing only soluble rec-PrP, was used for substrate preparation. The dialyzed proteins, with a concentration of around 20 µM given similar yields upon refolding, were diluted in conversion buffer (CB), consisting of 0.15M NaCl, 1% Triton-X-100 in PBS to which dextran sulfate sodium salt from *Leuconostoc spp*. with molecular weights ranging from 6500 to 10,000 (Sigma–Aldrich) was added to a final concentration of 0.5% (w/v). All substrates were aliquoted and stored at −80 °C until required.

### Protein Misfolding Shaking Amplification

To induce spontaneous misfolding of the recPrP variants and evaluate their misfolding proneness, a specific Protein Misfolding Shaking Amplification (PMSA) methodology, as described previously (36) was used. Briefly, substrates were disposed in eight 2 ml tubes with conical bottom and screw cap (Fisherbrand), four of them complemented with 100 mg of 1 mm diameter acid-washed glass beads (BioSpec Products, Inc.), and the other four with 100 mg of 0.1 mm diameter acid-washed glass beads (BioSpec Products, Inc.). All 8 tubes for each one of the substrates prepared were then submitted to PMSA at 39 °C using a Thermomixer (Eppendorf) with internal temperature control and shaking at 700 rpm continuously for 24 h. Four 24 h serial PMSA rounds were performed for each substrate, diluting 1:10 the PMSA product from the previous round into a new set of 8 tubes with freshly thawed substrate and their corresponding beads.

#### RecPrP^res^ detection

All PMSA products from the four serial rounds were transferred from the reaction tubes to 1.5 ml Eppendorf tubes and digested by adding proteinase K (PK) (Roche) at 25 µg/ml for 1 h at 42 °C in an oven (Nahita). Following digestion, samples were centrifuged at 19,000 × *g* at 4 °C for 15 min, the supernatant discarded, and the pellets resuspended and washed with at least 700 µl of PBS (Fisher Bioreagents). Then, samples were centrifuged for another 5 min at 19,000 × *g* and 4 °C, and the pellets were resuspended in 15 µl of loading buffer 4× (NuPage LDS, Invitrogen), previously diluted to 1× in PBS. PK-resistant PrP detection was performed through SDS-PAGE and Coomassie blue staining. PK-digested samples were boiled for 10 min at 100 °C and loaded onto 4–12% acrylamide gels (NuPAGE Midi gel, Invitrogen Life Technologies). The electrophoresis was run for 1 h and 20 min (10 min at 70 V, 10 min at 110 V, and 1 h at 150 V) and stained with BlueSafe (NZYTech) for 1 h at room temperature. To minimize variations from comparing samples run in different gels, the same amount of PMSA product was processed in all cases, and the different process times were strictly controlled, especially the staining time.

Additionally, different PMSA substrates (containing similar rec-PrP concentrations) were run along the digested PMSA products in the gels, serving as internal controls of the staining reagent to avoid dramatic differences between gels. All gels were scanned using the iBright™ CL750 Imaging System (Invitrogen) and analyzed using AlphaView software (version 3.4.0, ProteinSimple).

## Acknowledgments

We thank Albina Román-Castro and María José Pazos-Guldrís, from the University of Santiago de Compostela, for TEM imaging.

## Funded by

Spanish National Research Agency (AEI), Spanish Government, co-financed by the European Regional Development Fund-ERDF (grant numbers: PID2020-117465GB-I00 to JRR and PID2024-160022OB-I00 and PDC2025-165940-I00 to JC), Instituto de Salud Carlos III (ISCIII) through the project "AC23-2/00049" under the auspices of the EU Joint Programme for Neurodegenerative Diseases Research (JPND2023-108) - www.jpnd.eu, and Xunta de Galicia (grant ED431C 2024/11). RVC is a pre-doctoral fellow of the Xunta de Galicia. CIQUS is a “Singular Center” (grant ED431G 2023/03, partially funded by the EU ERDF. This work was also supported by EFA031/01 NEURO-COOP project, which is co-funded at 65% by the European Union through Programa Interreg VI-A España-Francia-Andorra (POCTEFA 2021-2027) granted to J.C. and H.E. Additionally, CIC bioGUNE currently holds a Severo Ochoa Excellence accreditation, CEX2021-001136-S also funded by Spanish National Research Agency.

## Supplementary Material

**Suppl. Fig. S1.**
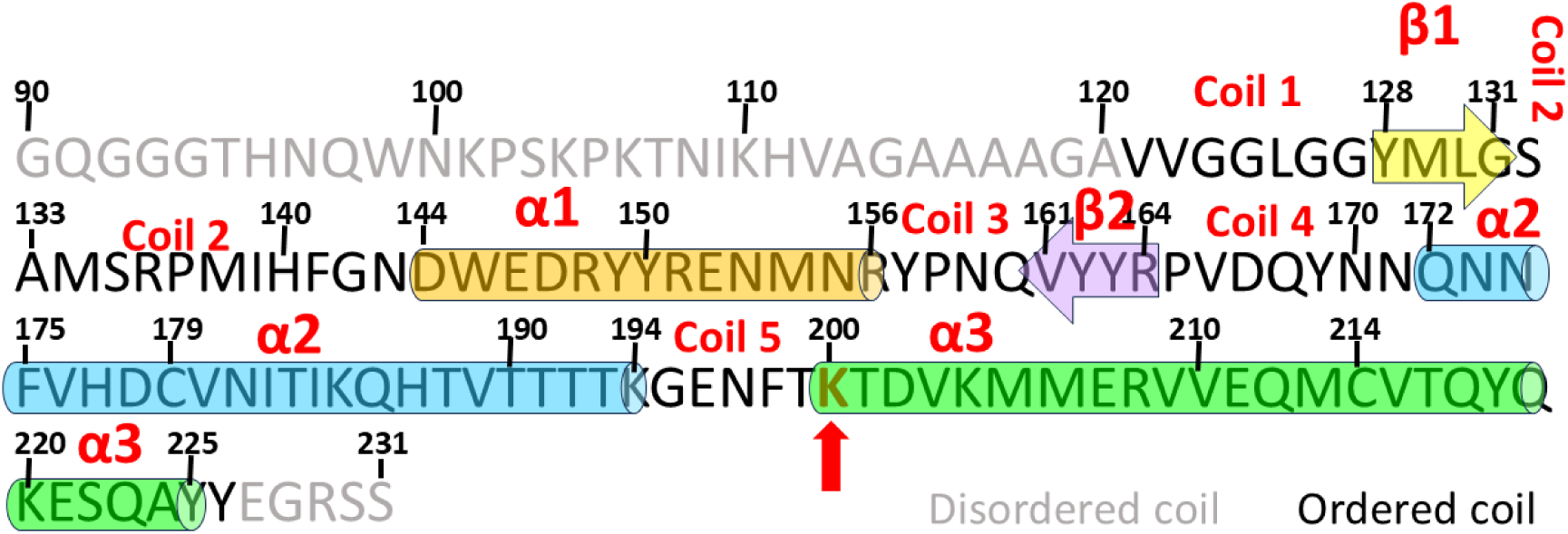
Cartoon showing the sequence of BVPrPE200K(90-231) with its secondary structure elements. The red arrow points to the E-to-K substitution. Secondary structure elements are based on those of wtBVPrP121-231 (14).

**Suppl. Fig. S2.**
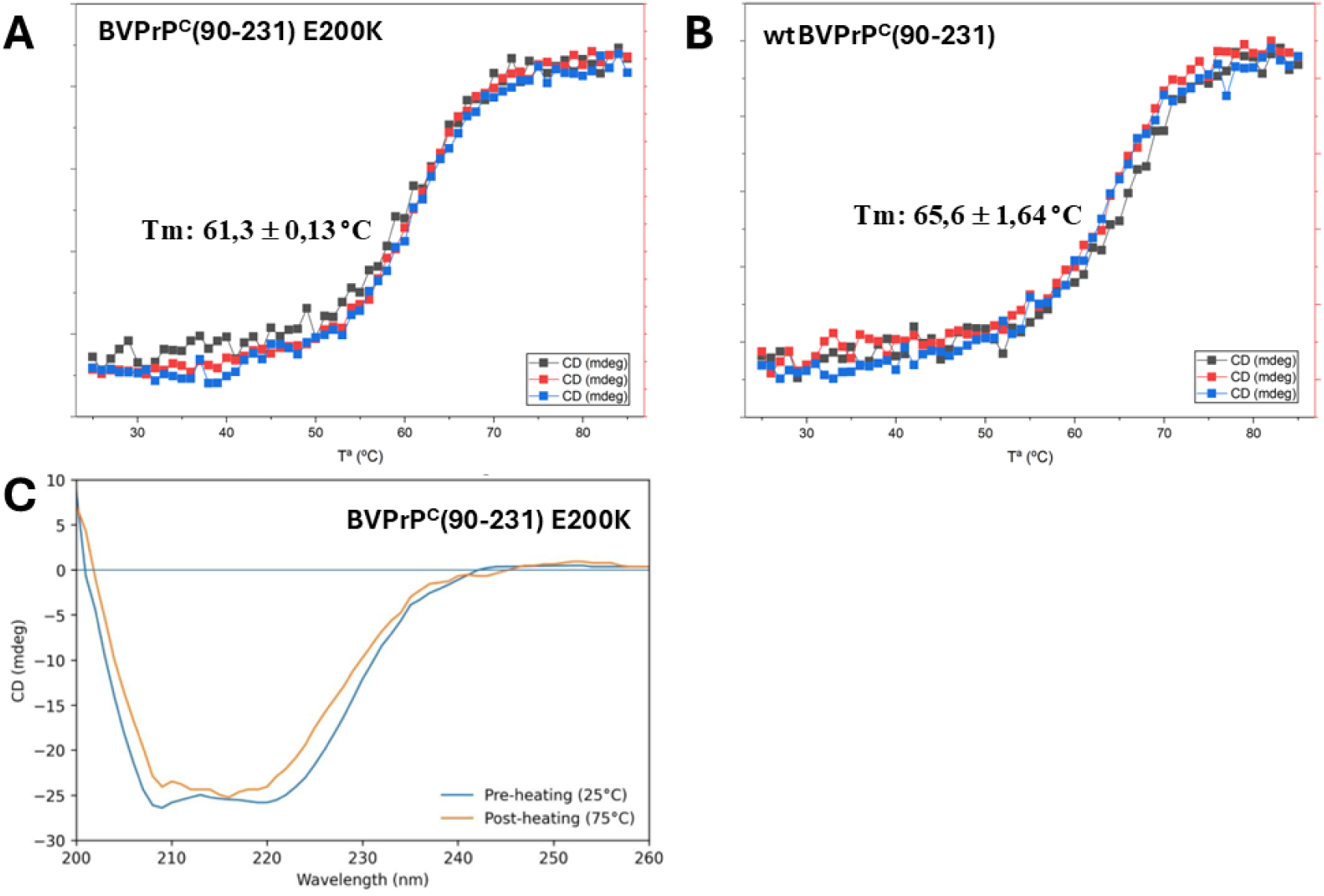
CD analysis of temperature-induced unfolding of recBVPrP^C^E200K(90-231). **A:** Unfolding curve evaluated as decrease in CD signal at 222 nm (α-helix content) over temperature of recBVPrP^C^E200K(90-231) (n =3 independent experiments); the datapoints were fitted to the Boltzmann sigmoidal function (Eq. 1) to calculate the Tm values. **B:** Same analysis of wtBVPrP^C^90-231) (n =3 independent experiments). **C:** Comparison of CD spectra of recBVPrP^C^E200K(90-231) at the beginning (25° C, fresh sample) and after heating at 75°C and cooling down to 25° C.

**Suppl. Fig. S3:**
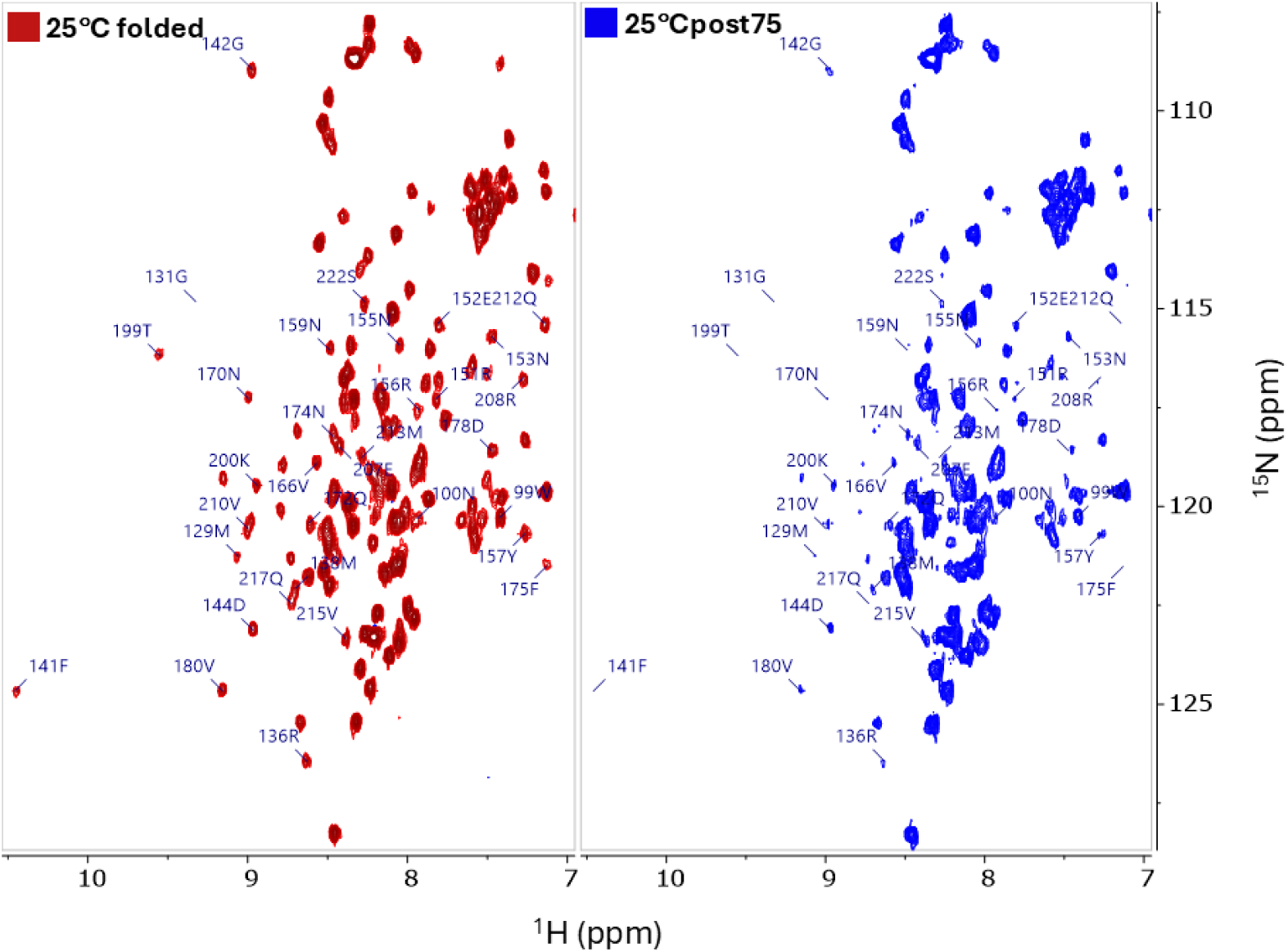
^1^H-^15^N HSQC spectra of BVPrP^C^E200K(90-231) with assignments (750 MHz). Comparison of spectra recorded at 25° C of the fresh sample (left panel, red) and the 25 °C post75 sample (right panel, blue). See main text for details. The protein dissolved in 10 mM sodium acetate buffer, pH 5, was supplemented with D_2_O (10%). The final protein concentration was 1.4 mg/ml.

**Suppl. Fig. S4.**
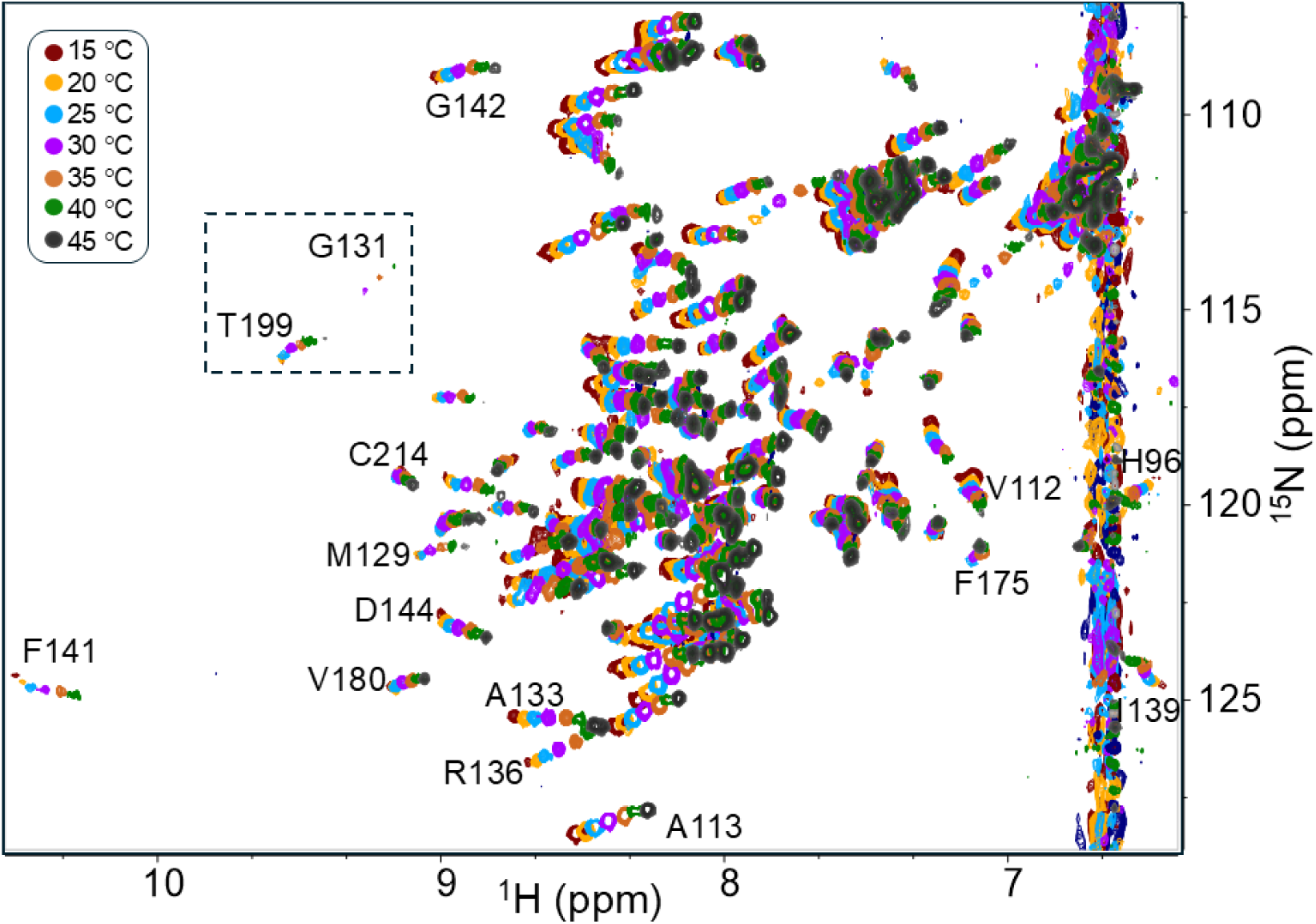
Temperature-induced chemical shift drift of recBVPrP^C^E200K(90-231) HSQC signals. The assignment of some of the peaks is shown. The inset highlights a signal (T199) with a linear drift with respect to temperature and another signal/one (G131) with a non-linear drift. Note also that some signals undergo a considerable change with temperature while others undergo a more modest one; this reflects in different Tc values.

**Suppl. Fig. S5.**
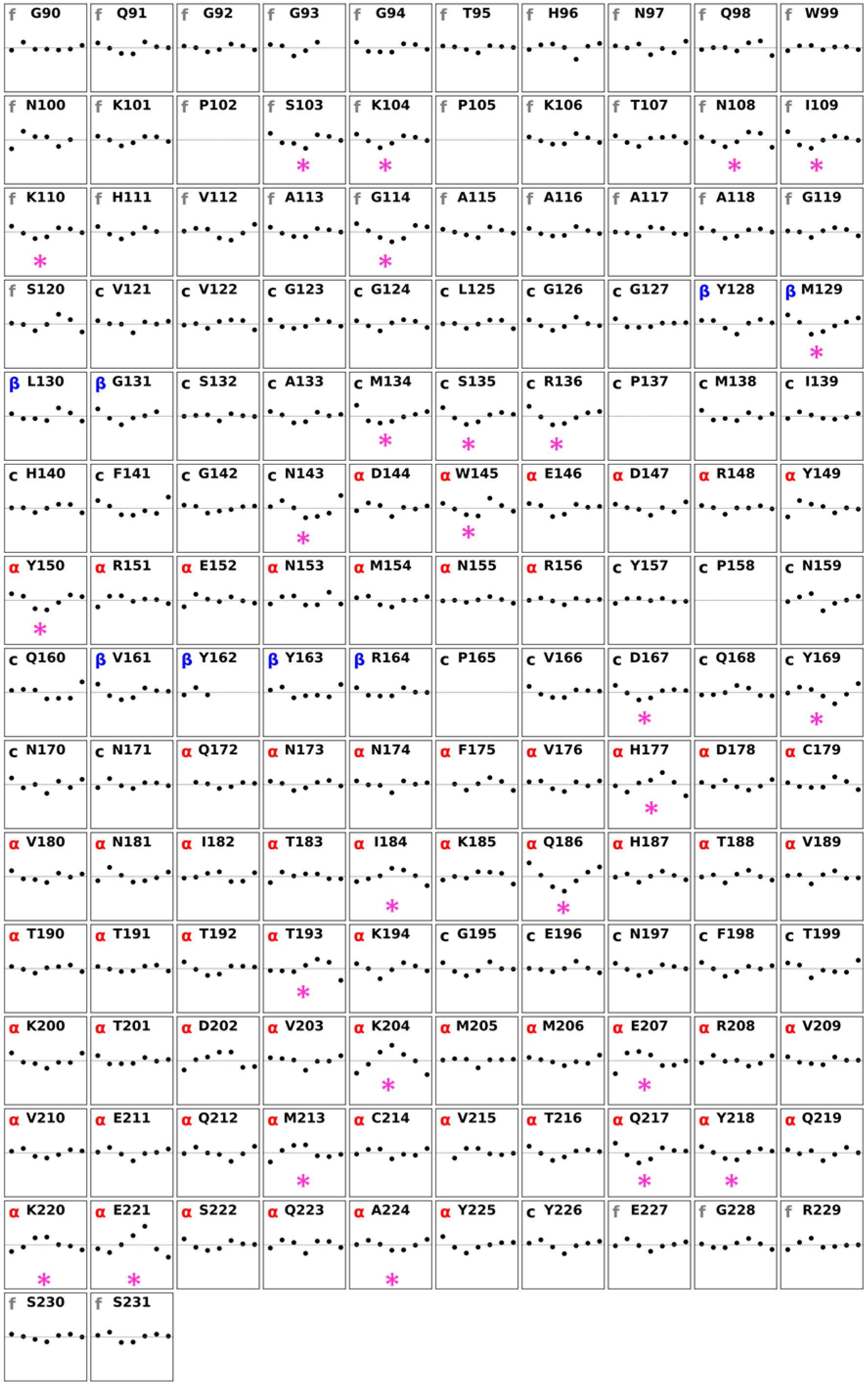
Plot of CCSD residual error for backbone amides of individual recBVPrP^C^E200K(90– 231) amino acid residues vs. temperature. The CCSD values for each residue were calculated by Eq. 2 through peak tracking across the series of VT N-HSQC spectra. They were fitted to a linear regression model with Eq. 3. Each plot represents the residual error (defined as CCSD - CCSD^calc^) versus temperature (31, 32). In each sub-plot the amino acid is shown, and the upper-left letter code refers to f=flexible (disordered, grey), α=helix (red), β=sheet (blue) or c=coil (black). The horizontal axis spans the temperature range 285–320 K, and the vertical axis ranges from −50 to +50 ppb. The magenta asterisk denotes a residue with non-linear variation of its chemical shifts with temperature.

**Suppl. Fig. S6.**
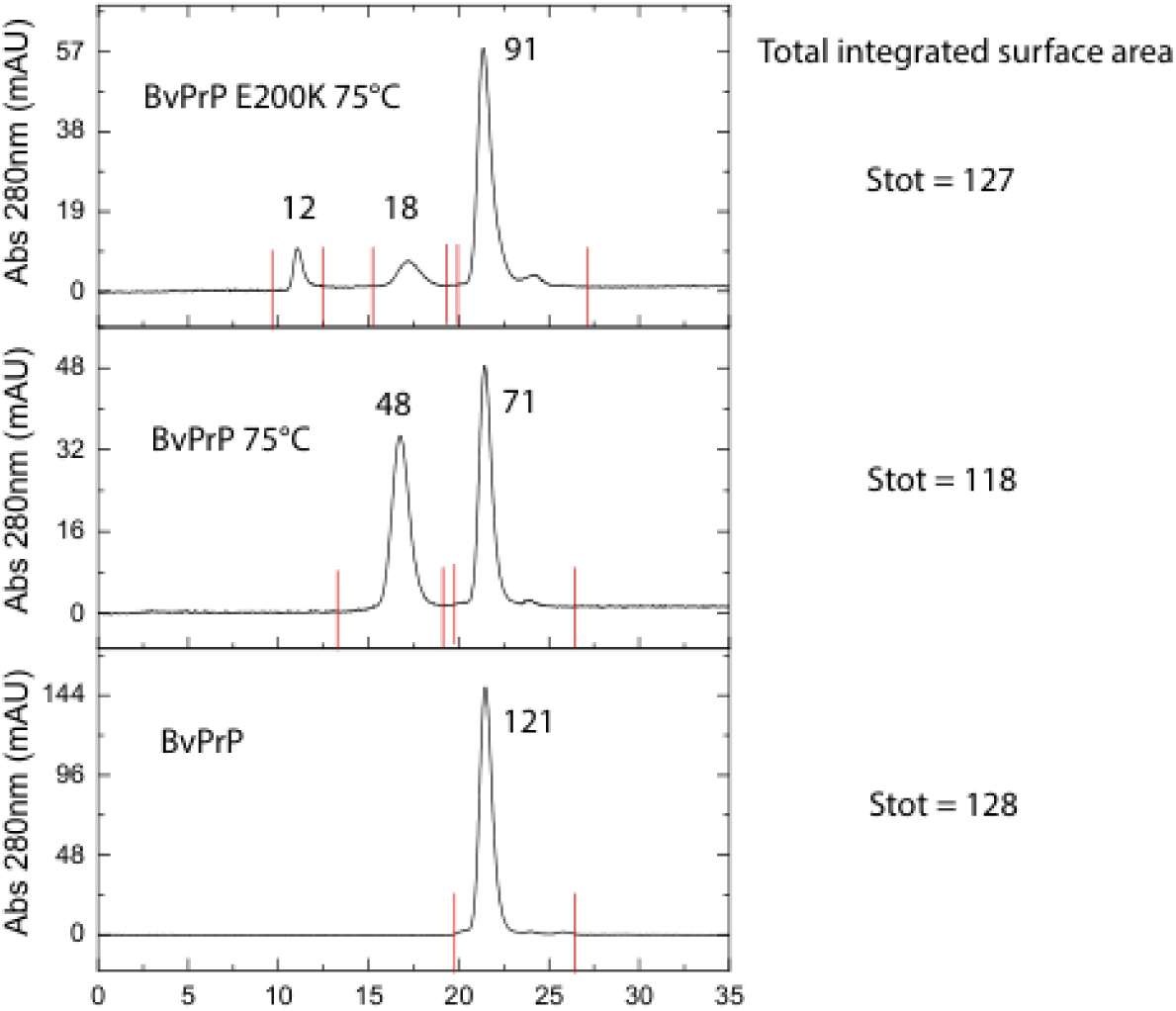
Relative amounts of oligomeric and monomeric species in post-75 °C wt and recBVPrP E200K (90-231) samples. Chromatograms are the same shown in Fig. 5A. Stot = total integrated area (arbitrary units). Numbers above each peak indicate the integrated areas of the peaks.

